# KCC1 Activation protects Mice from the Development of Experimental Cerebral Malaria

**DOI:** 10.1101/186262

**Authors:** Elinor Hortle, Lora Starrs, Fiona Brown, Stephen Jane, David Curtis, Brendan J. McMorran, Simon J. Foote, Gaetan Burgio

## Abstract

*Plasmodium falciparum* malaria causes half a million deaths per year, with up to 9% of this mortality caused by cerebral malaria (CM). One of the major processes contributing to the development of CM is an excess of host inflammatory cytokines. Recently K+ signaling has emerged as an important mediator of the inflammatory response to infection; we therefore investigated whether mice carrying an ENU induced activation of the electroneutral K+ channel KCC1 had an altered response to *Plasmodium berghei*. Here we show that Kcc1^M935K/M935K^ mice are protected from the development of experimental cerebral malaria, and that this protection is associated with an increased CD4+ T cells and TNF-α response. This is the first description of a K+ channel affecting the development of experimental cerebral malaria.

## Introduction

*Plasmodium falciparum* malaria is a major cause of mortality worldwide, causing an estimated 219 millions cases and 435,000 deaths in 2018^1^. One of the most severe and lethal complications of *P. falciparum* infection is the sudden onset of seizures and/or coma, known as cerebral malaria (CM). Its occurrence varies from region to region, with a case fatality rate as high as 9% of severe malaria cases in some areas ^2,3^. The causes of CM are not well understood, but hypotheses include the accumulation of parasitized red blood cells in the brain microvasculature, as well as imbalance in the pro- and anti-inflammatory responses to infection^4^.

In recent years, potassium (K+) signaling has emerged as an important mediator of the immune response to infection. Several studies have shown *in vitro* that functional outwardly rectifying K+ channels are necessary for macrophage activation and production of TNFα^5,6^, for activation of the NALP inflammasome^7^, for the activation of T helper cells, and the formation of T regulatory cells ^8–10^. The K+ content of the RBC also has a large effect on intra-erythrocytic *Plasmodium*. It has been shown that an outwardly directed K+ gradient is needed for normal parasite growth and maintenance of the parasite plasma membrane potential ^11–13^.

A mouse line expressing an activated form of K-Cl co-transporter type 1 (KCC1), discovered from a large scale ENU mutagenesis screen in mice, has recently been described^14^. The induced mutation – an M to K substitution at amino acid 935 of the protein – impairs phosphorylation of neighboring regulatory threonines, leading to over-activation of the transporter. The resulting increase in K+ efflux from the RBC causes *Kcc1^M935K^* mice to display microcytic anemia, with homozygous mutants showing a 21% decrease in Mean Corpuscular Volume (MCV), 8% decrease in total hemoglobin, and 21% increase in number of red cells. Mutant cells are also significantly less osmotically fragile^14^ indicating a dehydration of the red blood cells.

Here we use the *Kcc1^M935K^* mouse line to investigate the effect of increased host K+ efflux on susceptibility to malaria infection. When *Kcc1^M935K^* mice were infected with *Plasmodium berghei*, they showed protection from the development of experimental cerebral malaria (ECM), associated with a significant increase in CD4+ T cells and TNFα in the brain during infection, suggesting K+ efflux through KCC1 alters the inflammatory response to infection. This is the first description of a cation cotransporter affecting the development of ECM in mice.

## Results

### Kcc1^M935K^ mice are protected against P. berghei infection

Kcc1^M935K/M935K^ mice were inoculated with *P. berghei* to determine their resistance to parasitic infection. Cumulative survival and peripheral parasitemia were monitored daily over the course of infection. When mice were infected with 1×10^4^ *P. berghei* parasitized red cells, survival was significantly increased in the mutants, with 100% of homozygotes surviving past day 10 of infection, compared to 7% of WT females (P=0.0004; Figure 1A), and 11% of WT males (P < 0.0001; Figure1B). Significantly lower parasitemia was observed in both Kcc1^M935K^ females and males. In females, parasitemia was reduced by 63% on day 7, 48% on day 8, and 42% compared to WT, on day 9 post inoculation. Parasitemia in males was similarly reduced, by 66%, 41%, and 53% respectively (Figure 1A-D).

**Figure 1:**
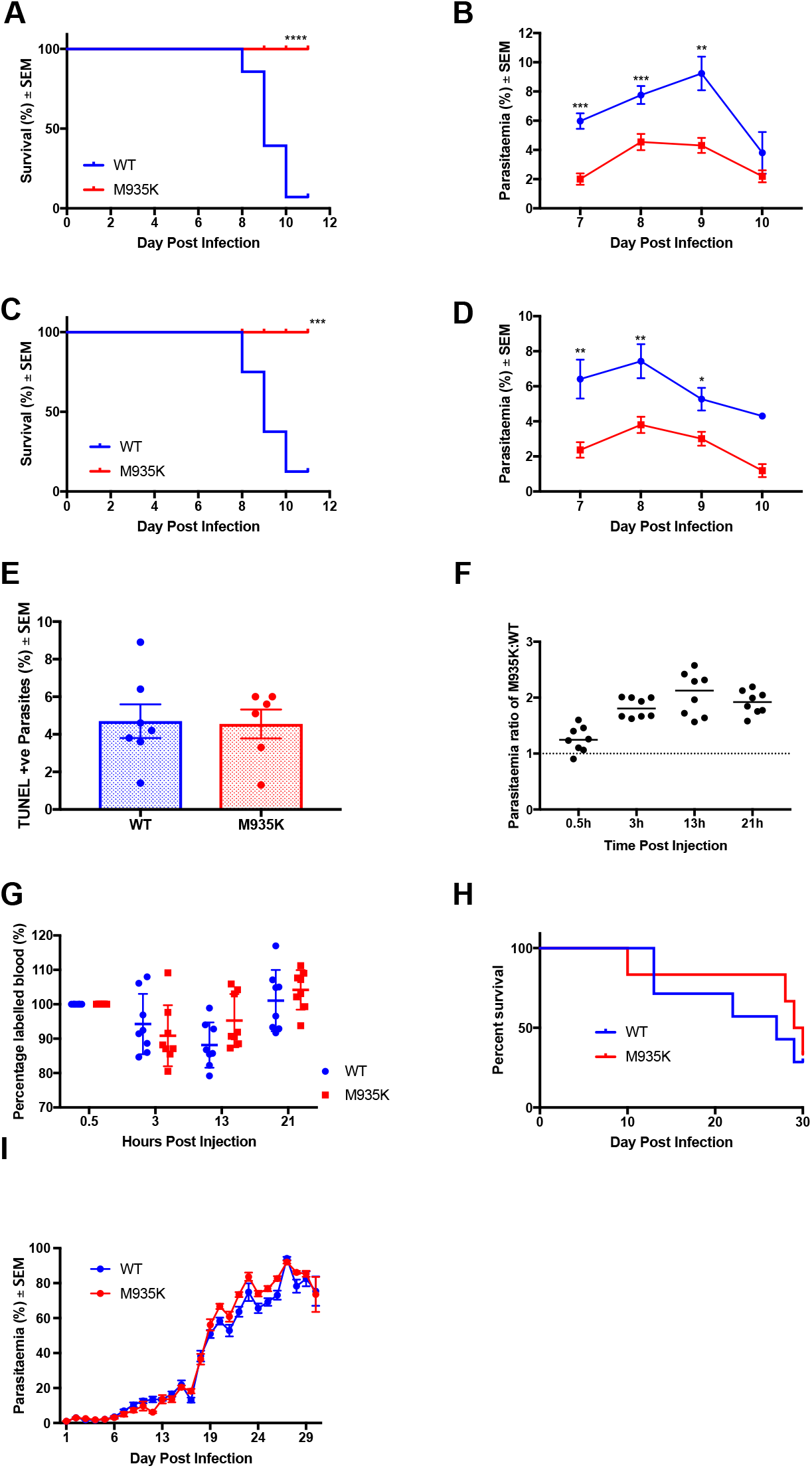
The Kcc1^M935K^ mutation causes resistance to *P. berghei*. **(A)** and **(B)** Cumulative survival and average ± SEM parasitemia for WT and Kcc1^M935K/M935K^ in male mice. WT n=28, WT Kcc1^M935K/M935K^ n=14. Combined results of two independent experiments. **(C)** and **(D)** Cumulative survival and average ± SEM parasitemia for WT and Kcc1^M935K/M935K^ in female mice. WT n=8, Kcc1^M935K/M935K^ n=10. Combined results of two independent experiments. *P<0.05, **P<0.01, ***P<0.001. P values calculated using Log rank test or the student’s T-test. **(E)** average ± SEM percentage of parasites which are TUNEL positive. **(F)** and **(G)** Average ± SEM fold change in parasitemia and percentage of remaining Kcc1^M935K/M935K^ labelled cells compared to WT labelled cells injected into the same *P. berghei* infected host (n=8) from 30 minutes to 21 hours post inoculation. **(H)** and **(I)** Cumulative survival and average ± SEM parasitemia for WT and Kcc1^M935K/M935K^ in female mice in challenge where WT did not develop CM. WT n=7, Kcc1^M935K/M935K^ n=5.

To determine if this reduction in parasitemia was caused by impairment in the parasite’s ability to invade Kcc1^M935K/M935K^ RBCs and survive within them, we conducted TUNEL staining of infected RBCs to detect fragmented nuclei in the parasites indicative of maturation arrest^15^, and an *in vivo* invasion and maturation assay as previously described^16^. No significant differences were observed in the TUNEL assay (Figure 1E) but a significant 2-fold increase in parasitic growth for Kcc1^M935K/M935K^ RBCs at 13 hours (P < 0.001) and 21 hours (P < 0.001) (Figure 1F), suggesting the Kcc1^M935K^ mutation does not affect parasite invasion but promotes intra-erythrocytic survival. To explain increased intra-erythrocytic survival but lower parasitemia for Kcc1^M935K/M935K^ over the course of infection (Figure 1B-D) we hypothesized that parasitic clearance was increased in the mutants. To address this postulate, we measured the proportion of labeled RBCs for WT and Kcc1^M935K/M935K^ RBCs across time from 30 minutes to 24 hours post inoculation and found no difference in the proportion of remaining Kcc1^M935K/M935K^ to WT labeled RBCs to WT indicative of no increase in parasitic systemic clearance and/or sequestration for Kcc1^M935K/M935K^ RBCs.

In the experiments described above, 90% WT mice succumbed between days 8 and 10 post infection. Death at this point in *P. berghei* infection is usually caused by experimental cerebral malaria (ECM), and unrelated to the level of peripheral parasitemia. Combined with our results from the TUNEL and invasion assays, this hinted that the Kcc1^M935K^ mutation may not affect parasite growth, but may rather provide more systemic protection from ECM. We therefore infected mice with a dose of *P. berghei* at which 50% of WT mice survived beyond day 10 of infection. In this experiment Kcc1^M935K/M935K^ had no difference in survival and no significant differences in parasitemia (Figure 1G-H). This suggests that parasite invasion or maturation is unlikely to be defective in Kcc1^M935K^ mutant mice.

### Kcc1^M935K^ promotes resistance to ECM

To determine if the WT mice were succumbing to *P. berghei* infection in our challenges resulted from as a result of ECM, mice were injected with *P. berghei* and symptoms of ECM were scored according to severity, from 0 (no symptoms) to 5 (death). Severe clinical symptoms were observed in WT mice, with most dying from seizures, whereas Kcc1^M935K/M935K^ remained asymptomatic for the length of the experiment (Figure 2A). One of the key hallmarks of cerebral malaria is breakdown of the blood brain barrier. Therefore, mice were injected intravenously with Evan’s Blue to assess blood brain barrier integrity. Infected WT mice showed an average of 14.5±2.9 grams of dye per gram of brain tissue, which was higher than the 8.1±1.4g/g observed in uninfected mice. Infected Kcc1^M935K/M935K^ more closely resembled uninfected mice, with 9.4 ± 1.3 g/g (Figure 2B-C). Although none of these differences was significant, the clear trend to higher staining in WT suggested a breakdown of the blood brain barrier that is not observed in Kcc1^M935K/M935K^. ECM is also associated with increased sequestration of infected RBCs in the microvasculature. We therefore assayed the relative amount of parasite sequestration in the brain and spleen by PCR of *P. berghei* 18S RNA. Kcc1^M935K/M935K^ showed a trend to decreased parasitemia in the brain, and increased parasitemia in the spleen (Figure 2D) indicating the difference in the ability of the parasite to cross the blood brain barrier. This trend was confirmed by measuring infected RBCs (iRBCs) in the brain by flow cytometry (Figure 2E). To further assess the postulate of an increase in sequestration of the parasitized RBCs in the brain tissue, we histologically assessed the presence of residual focal hemorrhages, oedema and neuropil changes in the brain tissue. At the pathological examination, the brain tissue appears normal for Kcc1^M935K/M935K^ or WT mice (Figure 3A-C), with no presence of residual hemorrhages, petechies or oedema. However we found the presence of hemozoin pigmentation with significant disruption of the neuropil accompanied with white blood cells infiltrations and uninfected RBC infiltrates in the infected WT mice at day 10 post inoculation (Figure 3B), whereas this was not observed for infected Kcc1^M935K/M935K^ mice (Figure 3D). Interestingly, pathological examination of the spleen indicates no architectural differences between infected and uninfected Kcc1^M935K/M935K^ and WT mice (Figure S1) but presence of hemozoin pigmentation in both strains indicates of splenic sequestration (Figure S1 C-D) and marked hyperproliferation of white blood cells in infected Kcc1^M935K/M935K^ mice (Figure S1-D) which was not evident in the spleen of infected WT mice (Figure S1-C). Together with the clinical scores and Evan’s blue staining, this suggests that WT mice are more likely to succumb to ECM due to a breakdown of the blood brain barrier and a moderate increase in iRBC population in the brain compared with Kcc1^M935K/M935K^ which are more likely to be resistant to the development of this condition, possibly through an increased sequestration of parasites in the spleen, which also aids in preventing iRBC localization to the blood brain barrier.

**Figure 2:**
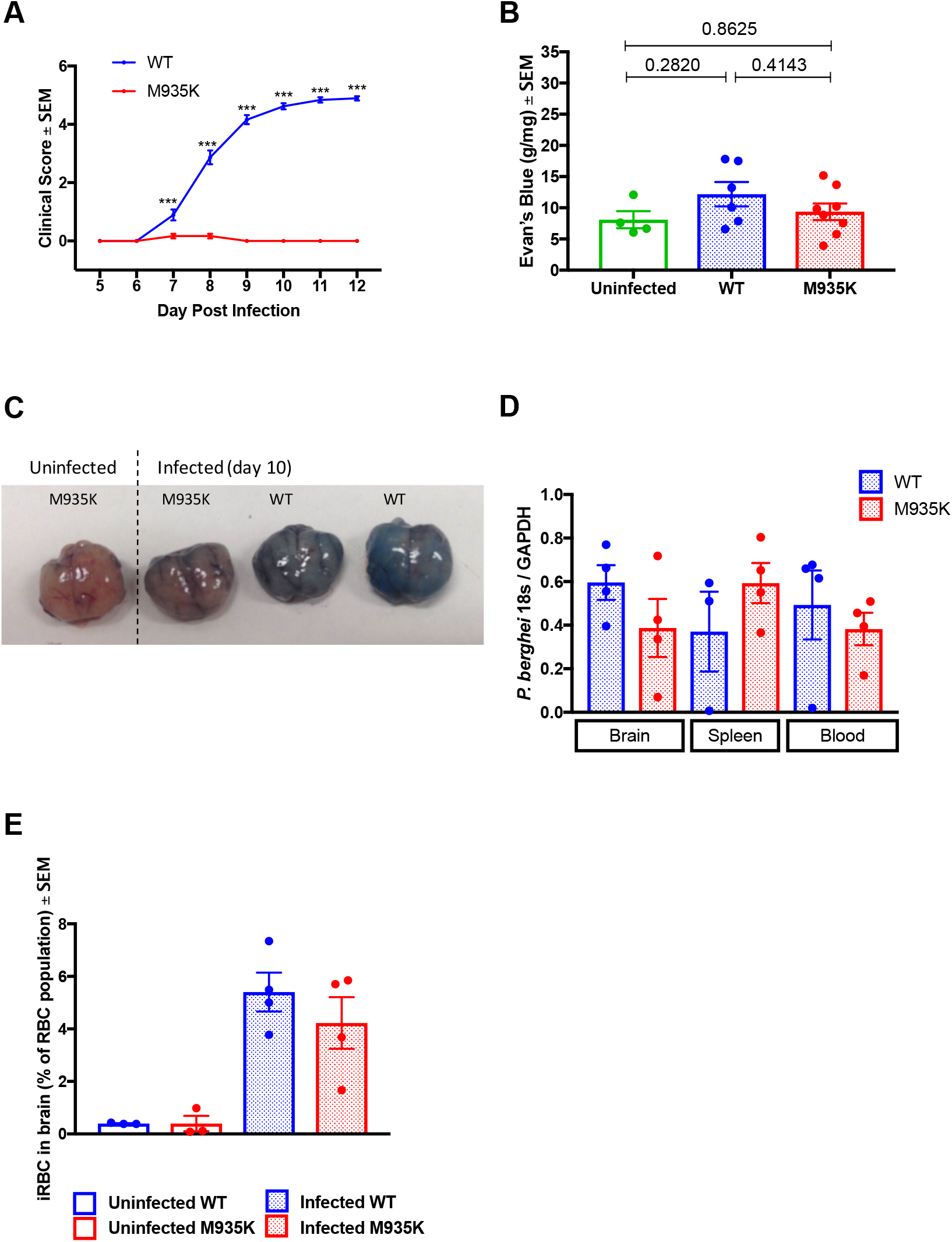
The Kcc1^M935K^ mutation causes resistance to cerebral malaria. **(A)** Clinical score for WT (n=37) and Kcc1^M935K/M935K^ (n=24) mice infected with *P. berghei* 0= no symptoms, 1= reduced movement, 2= rapid breathing/hunched posture, 3= ruffled fur/external bleeding, 4= fitting/coma, 5= death. **(B)** Amount of Evan’s Blue dye extracted from *P. berghei* infected Kcc1^M935K/M935K^ (n=8), WT (n=7) and uninfected (n=4) brains. **(C)** Representative brains dissected from mice injected with Evan’s Blue dye. **(D)** Relative amount of *P. berghei* 18s RNA normalized to mouse GAPDH in brain, spleen, and blood of infected Kcc1^M935K/M935K^ (n=4), WT (n=4). **(E)** Relative proportion of *P. berghei* infected Red Blood Cells in the brain from flow cytometry analysis. The population was gated on TER119 and Hoescht 33352 positivity. A comparison of the infected RBC (iRBC) in the brain of female WT and Kcc1^M935K/M935K^ mice, as an average percentage of total RBC in the brain ± SEM. Uninfected controls show low background staining as quantified using FACS. Uninfected WT and Kcc1^M935K/M935K^ n=3. Infected WT and Kcc1^M935K/M935K^ n=4. Values are average ± SEM. *P<0.05, **P<0.01, ***P<0.001. P values calculated using multiple T test with Two-stage linear step-up procedure of Benjamin, Krieger ad Yekutieli with Q=1% (A), or ordinary one-way ANOVA (B).

### Kcc1^M935K^ display an abnormal immune response to infection

It has been shown that depletion of CD4+ T cells, CD8+ T cells, and inflammatory monocytes can prevent the development of ECM ^17–19^. ECM resistance is also observed in mice with impaired thymic development of CD8+ T cells^20^. Since the parasites were present in the brain and KCC1 is expressed ubiquitously^21^, we postulated that the Kcc1^M935K^ mutation might cause alterations to some of these immune cell populations in the brain resulting in a stimulation of the immune response and impairment of parasite growth or increase clearance of the parasites. Therefore, the relative amounts of CD4+ and CD8+ T cells were measured by flow cytometry in the brain, blood, spleen, and thymus, both in uninfected mice, and the day the mice succumbed to ECM, where the inflammatory response is expected to be highest^22,23^. Consistent with the hypothesis that kCC1^M935K/M935K^ mice are resistant to ECM, differences in CD4+ and CD8+ T cells were observed in the brain (Figure 4 A-B), but not in the blood, spleen, or thymus (Figures 4C-D and S2). When uninfected, KCC1^M935K/M935K^ showed a 4-fold increase in the average number of CD4+ T cells in the brain compared to WT (Figure 3A). During infection, kCC1^M935K/M935K^ had a slight increase in the average amount of CD4+ T cells in the blood compared to WT (Figure 4C). Importantly, KCC1^M935K/M935K^ showed an 8-folds increase in CD4+ T cells in the brain during infection (P = 0.028), compared to the CD4+ T cells in infected WT, but only a 2-fold increase from their already higher baseline. The WT mice did not show a change in CD4+ T cells in the brain during infection (Figure 4A). Conversely, KCC1^M935K/M935K^ had a 2.5-fold lower in CD8+ T cell count compared to WT mice during infection (p = 0.028). There were no significant changes in CD8+ T cells in either genotype from their baseline when placed under infection in the brain (Figure 4B) or in the blood (Figure 4D).

**Figure 3:**
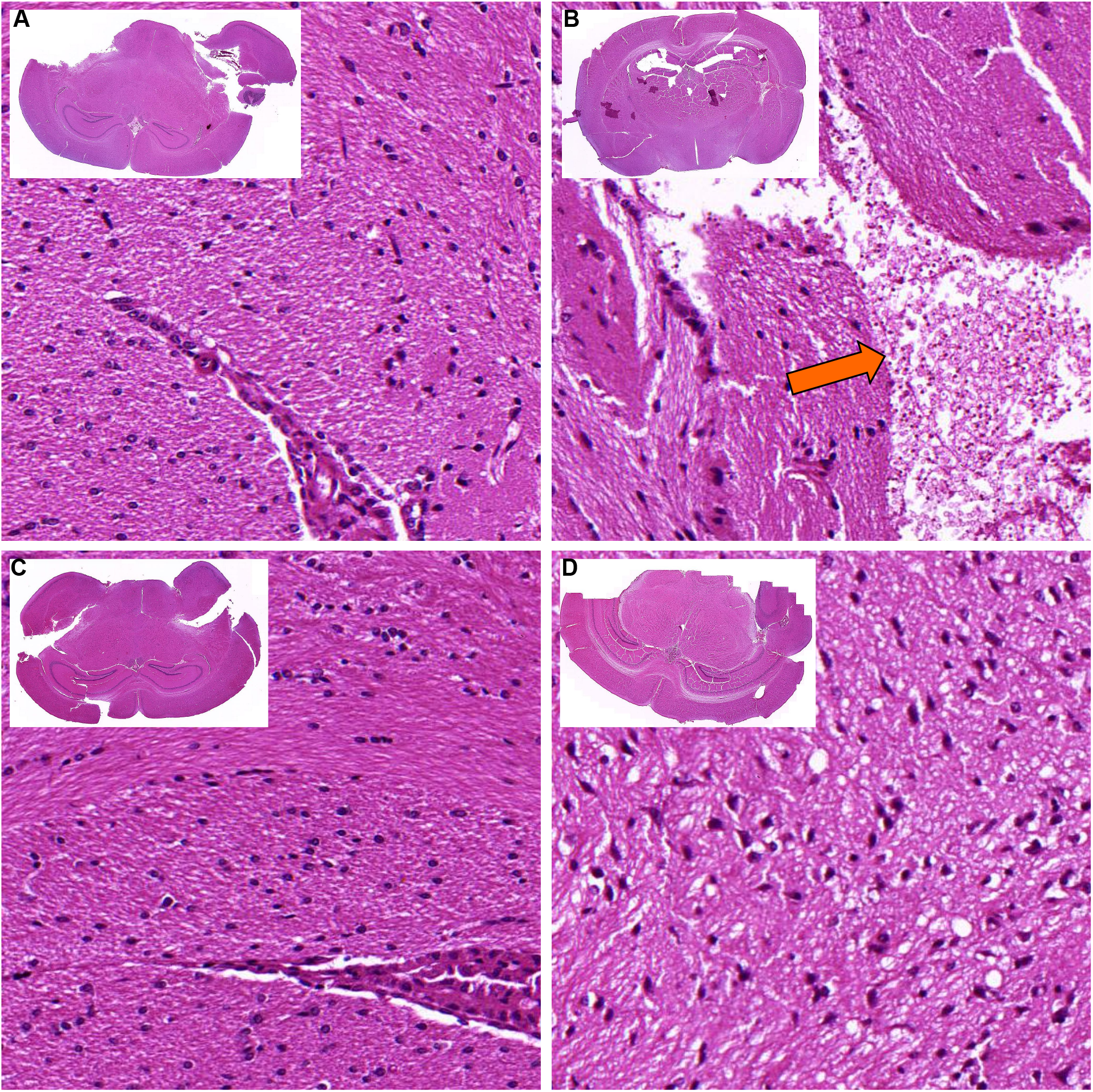
The Kcc1^M935K^ mutation protects the brain against parasitized RBCs. H&E stained brain sections from representative **(I)** uninfected WT, **(II)** infected WT, **(III)** uninfected Kcc1^M935K/M935K^, and **(IV)** infected Kcc1^M935K/M935K^ mice. The orange arrow indicates the presence of hemozoin pigmentation within the RBCs. Sections are 20x magnification, insets are 1.25× magnification.

**Figure 4:**
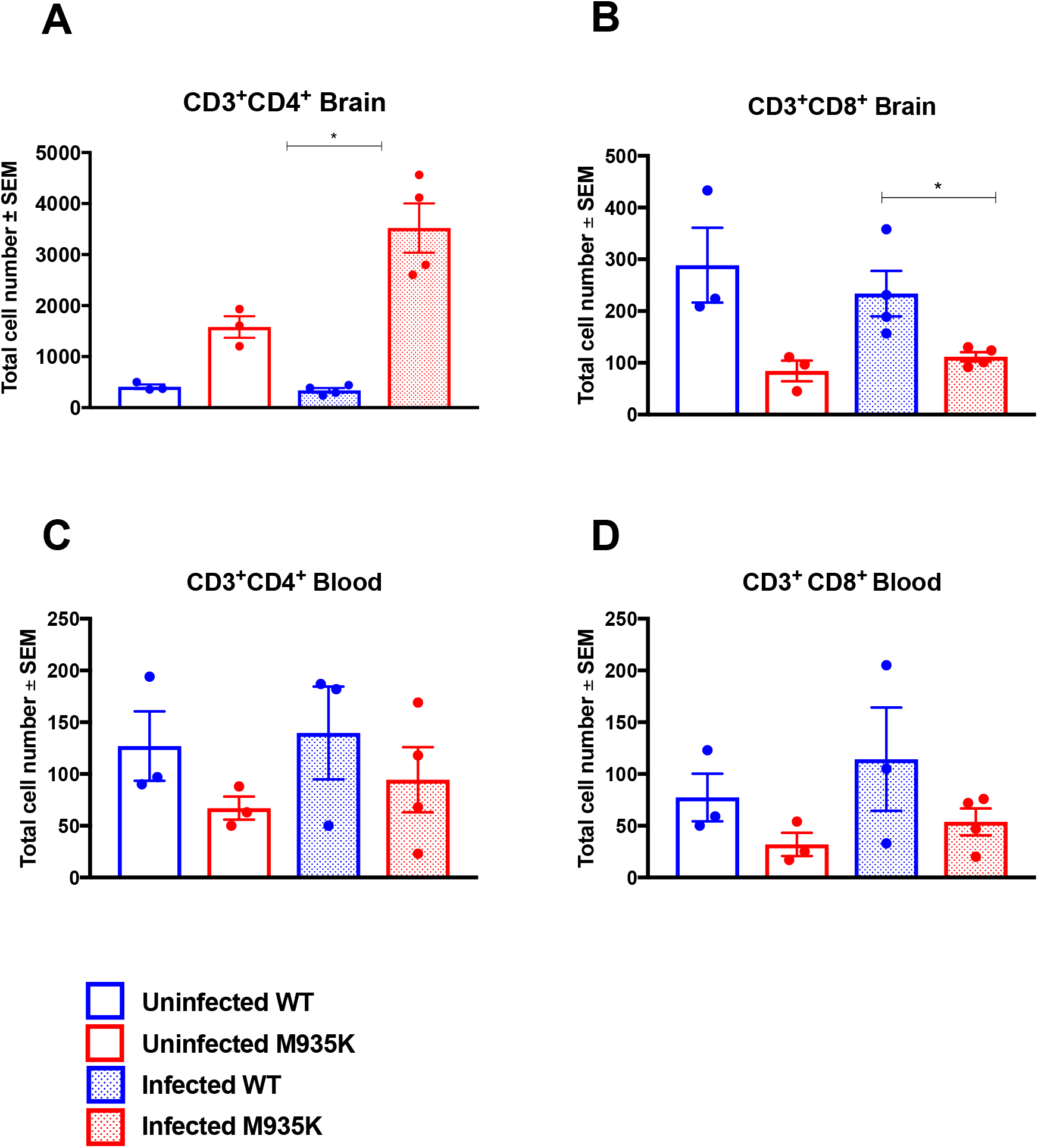
The Kcc1^M935K^ mutation alters CD4+ and CD8+ T Cell populations in the brain. Numbers of total cells in the brain that are **(A)** CD3+CD4+ and **(B)** CD3+CD8+ in uninfected WT and Kcc1^M935K/M935K^ (n=3), and *P. berghei* infected WT and Kcc1^M935K/M935K^ (n=4). Numbers of total cells in the blood that are **(C)** CD3+CD4+ and **(D)** CD3+CD8+ in uninfected WT and Kcc1^M935K/M935K^ (n=3), and *P. berghei* infected WT and Kcc1^M935K/M935K^ (n=4). Graph shows average ± SEM. *P<0.05, **P<0.01, ****P<0.0001. P values calculated using ordinary one way ANOVA.

One of the major host processes known to contribute to the development of cerebral malaria in *P. berghei* infection is an over-active inflammatory response. Both *in vivo* neutralisation of host molecules, and studies with knock-out mice have shown that cerebral malaria can be prevented by depletion of the pro-inflammatory cytokines IFN-γ ^24,25^ and TNFα^26^, and can be induced by depletion of the anti-inflammatory cytokine IL-10 ^27^ ECM resistance is also observed in mice with defective T cell dependent IFN-γ production ^20^. We therefore measured cytokine levels in infected mice in two ways: by ELISA in the brain and blood at a single time-point when all the WT mice succumbed to ECM, so all mice were sacrificed; and by CBA array in the plasma at several time-points over the first 10 days of infection.

At the single time-point, infected Kcc1^M935K/M935K^ showed a significant 1.7-fold increase in the average TNFα concentration in the brain (5723 pg/ml compared to 3329 pg/ml) and blood (2503 pg/ml compared to 1504 pg/ml) with a respective p-value of 0.0068 in the brain and 0.0134 in the blood (Figure 5A), as well as a trend to increased IL-1 *β* in the brain (Figure 5B). There were no differences in IFN-γ in either the brain or blood at this time-point (Figure 5C). By CBA array, Kcc1^M935K/M935K^ showed no difference in plasma cytokine levels early in infection; however, a 60% reduction in the amount of IFN-γ, and a 75% reduction in IL-6 were observed on day 9 of infection (391pg/ml compared to 969 pg/ml, and 2.4pg/ml compared to 10.3pg/ml respectively). This was followed by an 85% reduction in the amount of IL-10 on day 10 of infection (an average of 8pg/ml in Kcc1^M935K^ compared to 55pg/ml in WT) (Figure S3). A slight increase in the amount of TNFα (P=0.040) was also observed on day 7 (Figure S3). Together this indicates a possible protective effect of the Kcc1^M935K/M935K^ mice against ECM by altering the balance of IFN-γ and TNFα responses to infection.

**Figure 5:**
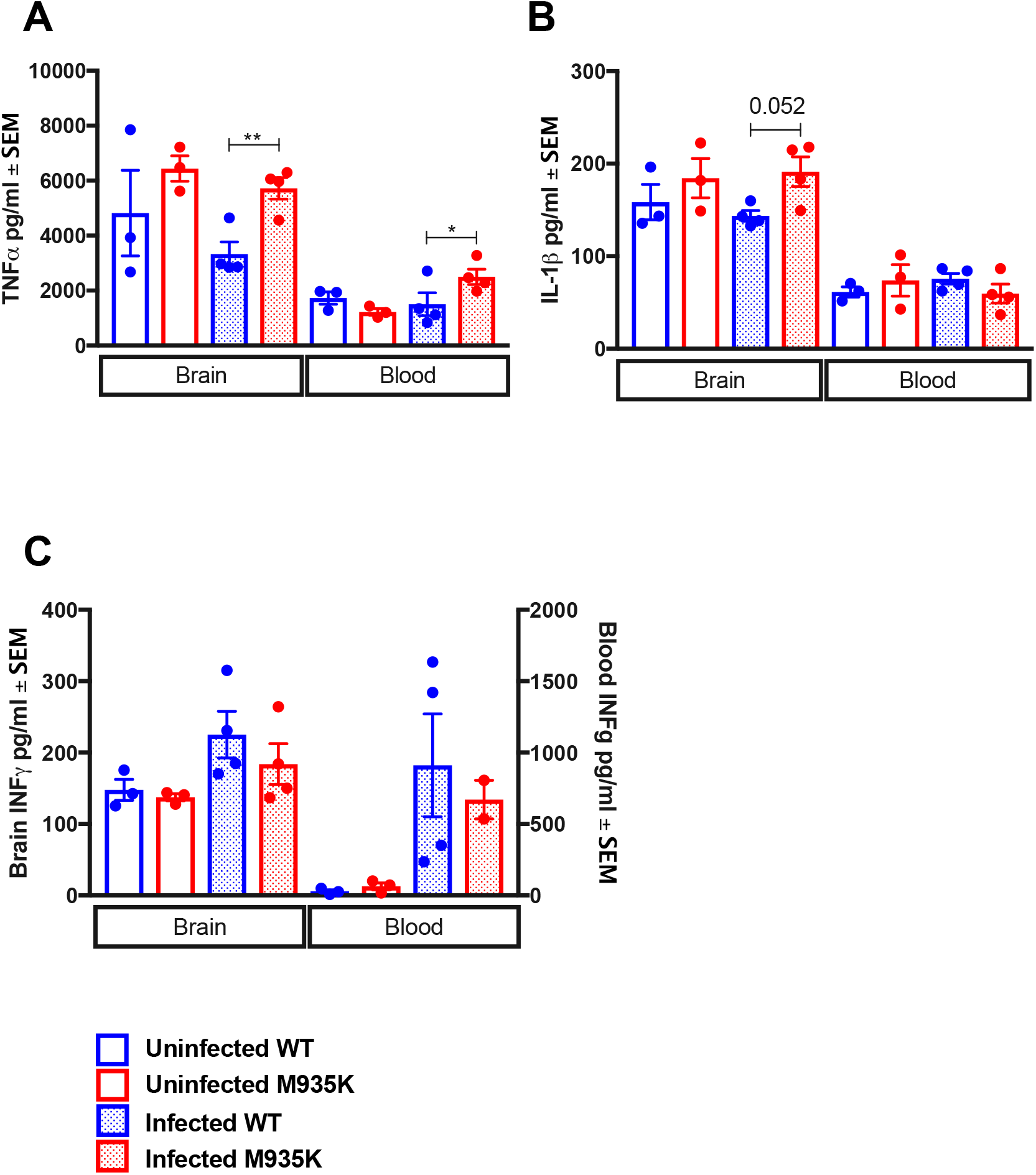
The Kcc1^M935K^ mutation increases pro-inflammatory cytokines in the brain. Average ± SEM amount of **(A)** TNF-α **(B)** IL-1β, and **(C)** INF-γ as measured by ELISA, in the brain and blood of uninfected WT and Kcc1^M9^ ^35K/M935K^ (n=3), and *P. berghei* infected WT and Kcc1^M935K/M935K^ (n=4). *P<0.05, **P<0.01 P values calculated using student’s T-test.

## Discussion

This study provides the first evidence that host KCC1 plays a role in malaria resistance. It shows that over-activation of the transporter is likely to provide protection to experimental cerebral malaria (ECM) in *P. berghei* infection. This is the first description of a mutation in a cation transporter that has an effect on ECM; other previously discovered genes have directly involved host cytokines, antigen presentation ^28,29^, or erythrocyte membrane proteins ^30–32^.

Despite the fact that Kcc1^M935K/M935K^ showed significantly lower parasitemia than WT during the first 10 days of infection, the mutation did not appear to have a cell autonomous effect on parasite invasion and survival within the RBC. One possible explanation for this observation is suggested by the fact that *P. berghei* infected RBCs have the ability to cytoadhere to the endothelium of blood vessels. Lower levels of sequestration in the mutants would leave more late stage parasites vulnerable to splenic clearance, and therefore result in reduced parasite burden^33^. Reduced sequestration would also be consistent with protection from cerebral malaria, as infected cells would be less likely to adhere within the microvasculature of the brain. Our results from *P. berghei* 18S rRNA and histology examination support this hypothesis, showing trends to reduced parasite burden in the brain and increased burden in the spleen but were not significant due to the genetic variation amongst mice.

Our results show increased CD4+ and decreased CD8+ T cells in the brains of infected KCC1^M935K/M935K^ compared with infected WT mice This corroborates previous reports showing CD8+ are the major mediators of ECM^17^, although it remains unclear from these experiments why mutants would have altered T cell populations in the brain even when uninfected. KCC1 is expressed on a wide range of immune cells, and may greatly affect their function. Previous studies have shown that K+ efflux can alter cellular cytokine production^5–7^; can increase assembly of the NALP inflammasome in response to pathogen associated proteins^7^; and is essential for macrophage migration^34^. All of these may contribute to both T cell migration, and the increased IL-1β and TNF-α observed in KCC1^M935K^ mice.

The increase of pro-inflammatory cytokines in the brains of KCC1^M935K^ mice was surprising, as increased TNF-α have previously been associated with more severe CM (reviewed in ^35,36^). However, it has been shown that neither TNF-α knock-out, or neutralization with antibodies, is sufficient to prevent CM ^37,38^, suggesting that the soluble cytokine is not itself causative of the condition. Although we did not find any significant differences in IFN-γ in the brains of KCC1^M935K^ mice at our single time point, we did observe a significant reduction in the plasma two days later than the significant increase in TNF-α. Interestingly, both IFN-γ knock-out and neutralization with antibodies, does protect against ECM (reviewed in ^39^). It may therefore be that the KCC1^M935K^ mutation does not protect by modulation of any one cytokine, but by a better ability to quickly reduce inflammation after its initial peak.

Here we have shown that activation of KCC1 is likely to provide protection to *P. berghei* by preventing the development of experimental cerebral malaria (ECM). This is the first description of a mutation in a transporter that has an effect on ECM. Previous studies have shown that pharmacological activation of KCC channels is achievable^40,41^, therefore future research into KCC1 activation may provide novel treatments for cerebral malaria.

## Methods

### Animals

Mice were bred under specific pathogen free conditions. All procedures conformed to the National Health and Medical Research Council (NHMRC) code of practice. All mouse procedures have been approved by the Australian National University Animal Experimentation Ethics Committee (AEEC A2014/054). The Kcc1^M935K^ mutation is carried on a mixed BALB/c and C57BL/6 background^14^. These two mouse strains differ in their susceptibility to *P. berghei*, and this introduced a greater amount of variability into results than is usually observed. Therefore, WT x WT and Kcc1^M935K/M935K^ x Kcc1^M935K/M935K^ breeding pairs were maintained. To exclude the possibility that the resistance phenotype was due to the mixed background, and carried by chance in mutant breeding pairs, Kcc1^M935K^ was periodically crossed back to WT, and new WT × WT and Kcc1^M935K/M935K^ × Kcc1^M935K/M935K^ pairs established from the progeny.

### Infections

Experiments used the rodent parasite *P. berghei ANKA*. Parasite stocks were prepared from passage through resistant SJL/J mice, as described previously^42^. Experimental mice were infected intraperitoneally at a dose of 1×10^4^ or 1×10^5^ parasitised RBC per mouse. Blood stage parasitemia was determined by counting thin smears from tail blood stained in 10% Giemsa solution. A least 300 cells were counted per slide.

### Histology

Uninfected, *P. berghei* infected mice and control were humanely euthanized. The brain was transcardially perfused by with 10 ml of 0.1M ice cold phosphate buffered saline solution (PBS) followed by 10 ml of ice cold 4% paraformaldehyde (PFA) and fixed into 70% ethanol. Brains and spleens from 5 mice from each group were serially sectioned and stained with Hematoxilyn and Eosin (H&E). Brain and spleen sections were independently examined from 2 different pathologists.

Thin tail smears from *P. berghei* infected mice were fixed in 100% MeOH, and stained with an APO-BrdU TUNEL assay kit according to the manufacturer’s instructions (Invitrogen, Carlsbad, CA). Slides were examined on an upright epifluorescence microscope (ZIESS) 600x magnification. 10 fields of view were counted for each slide.

### Evans Blue

*P. berghei* infected mice, and uninfected controls, were injected via IV with 200μl 1% Evans Blue/PBS solution. 1hr post injection, mice were sacrificed and their brains collected and weighed. Brains were placed in 2ml 10% neutral buffered formalin at room temperature for 48hrs to extract dye. 200μl of formalin from each brain was then collected and absorbance measured at 620nm. Amount of Evans blue extracted per gram brain tissue was calculated using a standard curve ranging from 40μg/ml to 0μg/ml. Injections were carried out on the day of infection that the first mouse died.

### Clinical Score

Mice were monitored three times daily, and given a score from 0 to 5 based on the type and severity of their symptoms. ‘0’ indicated no symptoms; ‘1’ reduced or languid movement; ‘2’ rapid breathing and/or hunched posture; ‘3’ ruffled fur, dehydration and/or external bleeding; ‘4’ fitting and/or coma; ‘5’ death. Mice were considered comatose if they were unable to right themselves after being placed on their side. The highest score recorded for each mouse on each day was used to generate daily averages.

### Cytokines

Peripheral blood was taken either by cardiac puncture or mandibular bleed in a microcentrifuge tube coated with anticoagulants and centrifuged for 4 minutes at 11,000xg. Plasma was then taken into a separate tube and stored at −20°C until needed. Cytokine analysis was conducted on un-diluted plasma using a CBA Mouse Th1/Th2/Th17 Cytokine Kit to the maker’s instructions (BD biosciences).

### Lymphocyte and Infected Red Blood Cell Analysis

Peripheral blood was taken either by cardiac puncture or mandibular bleed, and lymphocytes were isolated on Ficoll-Paque™ according to the maker’s instructions. Lymphocytes were then incubated with Fc-block in MT-FACS for 10 minutes at 4°C.

Both spleen and thymus were prepared for flow cytometry using the same method. ½ of each organ was passed through a 70μm BD Falcon Cell Strainer with 0.5 ml of MT-FACS buffer, and then centrifuged at 300xg for 5 minutes at 4°C. The supernatant was removed and the pellet re-suspended in 5 ml cold MT-FACS buffer. A 200μl aliquot of this suspension was then incubated with 0.8μl Fc-block.

Blood, spleen and thymus samples were then stained with CD4-PacificBlue, CD11-PE, CD8-FITC, CD25-PECy7, CD19-PERCPCy5.5 and CD3-APC-Cy7, then fixed with 300μl of MT-PBS containing 1% formalin and 1% BSA for 10 minutes at 4°C. Cells were then permeabilised with MT-PBS containing 0.1% saponin and 1%BSA at 4°C for 20 minutes. Finally, cells were stained with FoxP3-APC. Samples were acquired using a BD FACSAria™ II flow cytometer, and analysed using BD FACSDiva™ software (BD Biosciences).

For comparative analysis of CD3+CD4+ and CD3+CD8+ brain lymphocyte populations using flow cytometry, the brains of WT and mutant mice were harvested when WT mice succumbed to ECM. Briefly, the entire brain was passed through a 70μm BD Falcon Cell Strainer and collected post-straining in 1.5ml of PBS and then centrifuged at 500xg for 5 minutes at 4°C. The supernatant from this was used for ELISA (methods below). The cell pellet was then topped up to a total volume of 400μl with PBS, and split as 300μl for FACs staining, and 100μl for PCR (methods below). The samples were passed through a 70μm BD Falcon Cell Strainer again and then centrifuged at 1500xg for 5 minutes at 4°C, prior to washing once with PBS; 10μl of this pellet was then removed for infected RBC (iRBC) analysis. The remaining 40μl cell pellet was blocked with 5μl Fc block for 10 minutes at 4°C. The pellet was then washed twice with 200μl MTRC, and incubated with CD3-BV605, CD8-FITC and CD4-APC antibodies. Samples were acquired at 2.5×10^6^ cells per sample, using a BD LSRFortessa™, and analysed using BD FlowJo™ software.

The brain cell pellet previously removed for iRBC analysis, was incubated with TER-119-PE-Cy7 antibody, with Hoechst 33342, and JC-1 dyes in MTRC, and 2.5×10^6^ cells were acquired per sample using a BD LSRFortessa^™^, and analysed using BD FlowJo™ software.

### 18s PCR from Brain, Spleen and Blood

Brain samples were processed as described above. The spleen was also passed through a 70μm BD Falcon Cell Strainer and resuspended to a total volume of 1.5ml in PBS.

These cells were then centrifuged at 1500xg for 2 minutes at 4°C, and the supernatant was collected for ELISA (methods below). The pellet was resuspended to a total of 800μl and split in two, with 400μl taken for PCR, and 400μl snap frozen. Peripheral blood was collected using a cardiac puncture, and centrifuged at 1500xg 10 minutes at 4°C. The supernatant was taken for ELISA (below) and the pellet was lysed using two volumes of 0.15% saponin in PBS for 30 minutes at 37°C. Brain and spleen samples were processed similarly in order to successfully lyse any iRBC. The samples were then centrifuged at 10 000xg for 10 minutes at 4°C, and the pellet was washed three times with ice cold PBS. Following this, all samples were processed using the Qiagen DNeasy Blood and Tissue kit with following the manufacturer’s instructions. The quality and yield of DNA was quantified using a NanoDrop spectrophotometer. 200ng of each sample was set up in a PCR reaction with 1× MyTaq™ Mix and 0.4 μM of GAPDH primers (Forward: GATGCCCCCATGTTTGT; Reverse: TGGGAGTTGCTGTTGAAG), or 0.4μM of 18s primers (Forward: CAGACCTGTTGTTGCCTTAAAC; Reverse: GCTTGCGGCTTAATTTGACTC). The reaction parameters for GAPDH were as follows, 95°C for 5 minutes before 35 cycles of 95°C 30 seconds, 50°C 30 seconds, 72°C 1 minute, and a final extension at 72°C 2minutes. Since the genome of *P. berghei* is AT rich, the PCR protocol for 18s was 95°C 5 minutes, before 35 cycles of 95°C for 30 seconds, 50°C for 1 minute, 68°C for 2 minutes, and a final extension of 68°C for 5 minutes. These reactions were then eletrophoresed on a 1% agarose gel, and imaged using a Bio-Rad Gel Doc™ XR+ Gel Documentation System. Densitometry analysis was conducted on these bands using ImageJ software, and standardized to GAPDH densitometry analysis.

### ELISA from brain and blood

Samples were processed as described above, with the supernatants removed and saved for ELISA. ELISAs were conducted following the manufacturers’ protocols, with the IL-1b and IFN-y (ELISAKIT.com, EK-0033 and EK-0002, respectively) and TNF-a (ThermoFisher Scientific KMC4022). Samples were diluted 1 in 5 and 1 in 10 for IL-1b; 1 in 4 and 1 in 8 for IFN-y, and 1 in 10 and 1 in 20 for TNF-α. The pdata was collected using a Tecan M200 plate reader.

### *In-vivo* Invasion Assay

Blood from Mutant and WT uninfected mice was collected by cardiac puncture. 1800 *μ* l of blood was collected and pooled for each genotype, then halved and stained with either NHS-Atto 633 (1 *μ*l/100*μ*l) or sulfobiotin-LC-NHS-Biotin (1*μ*l/100*μ*l of 25mg/ml in DMF). Cells were then incubated at RT for 30 minutes, and washed twice in MT-PBS. Stained cells were combined in equal proportions to achieve the following combinations:

1) WT-Biotin + Mutant-Atto 2) WT-Atto + Mutant Biotin

Combined cells were then resuspended in 2ml MT-PBS, and injected intravenously into 4 WT *P. berghei* infected mice at 1-5% parasitemia, plus 1 uninfected control, were injected with 200 μ l dye combination 1; the same numbers of mice were injected with 200 μl dye combination 2. Injections were carried out when parasites were undergoing schizogeny, at ~1am.

30 minutes post injection, 1 *μ* l tail blood was collected and stained for 30 minutes at 4°C in 50 *μ*l MT-PBS containing 0.25 *μ* l CD45-APC-Cy7, 0.25 μ l CD71-PE-Cy5, 0.5 μ l Step-PE-Cy7. Next, 400 μ l MTPBS containing 0.5 *μ* l Hoechst 33342 and 1 *μ* l 800 μ g/ml Thiazole orange was added, and cells were incubated for a further 5 minutes at 4°C. Stained cells were then centrifuged at 750xg for 3 minutes, re-suspended in 700μl MT-PBS, and analyzed on a BD Fortessa Flow Cytometer. 2×10^6^ cells were collected for each sample, and data was analysed using FlowJo (FlowJo, LLC, Oregon, USA).

### Statistical analysis

P values were determined using Log-rank test, Welch’s T test or ordinary one-or two-way ANOVA where appropriate. Statistics on the clinical score (Figure 2A) was determined using two-stage linear step-up procedure of Benjamin, Krieger ad Yekutieli with Q=1%.

## Acknowledgement

We would like to acknowledge Shelley Lampkin and Australian Phenomics Facility (APF) for the maintenance of the mouse colonies. This study was funded by the National Health and Medical Research Council of Australia (Program Grant 490037, and Project Grants 605524 and APP1047090), the National Collaborative Research Infrastructure Strategy (NCRIS), the Education Investment Fund from the Department of Education and Training, the Australian Phenomics Network, Howard Hughes Medical Institute and the Bill and Melinda Gates Foundation. This study utilizes the Australian Phenomics Network Histopathological and Organ Pathology service, University of Melborne. We finally would like to thank Dr John Finnie, University of Adelaide, Australia and Professor Catriona A. McLean from the University of Melbourne, Australia for their assistance. We finally wish to acknowledge two anonymous reviewers for the careful reading of the manuscript and their helpful suggestions to improve the quality of this work.

## Author Contribution Statement

E.J.H, B.J.M, S.F.J and G.B designed and planed the experimental work. E.J.H, L.S and F.C.B performed the research. E.J.H, S.M.J, D.J.C. B.J.M, S.F.J and G.B interpreted and analyzed the data. E.J.H, L.S. and G.B performed the statistical analysis. E.J.H, L.S and G.B wrote the manuscript. All authors reviewed the manuscript.

## Supplementary material

**Figure S1:**
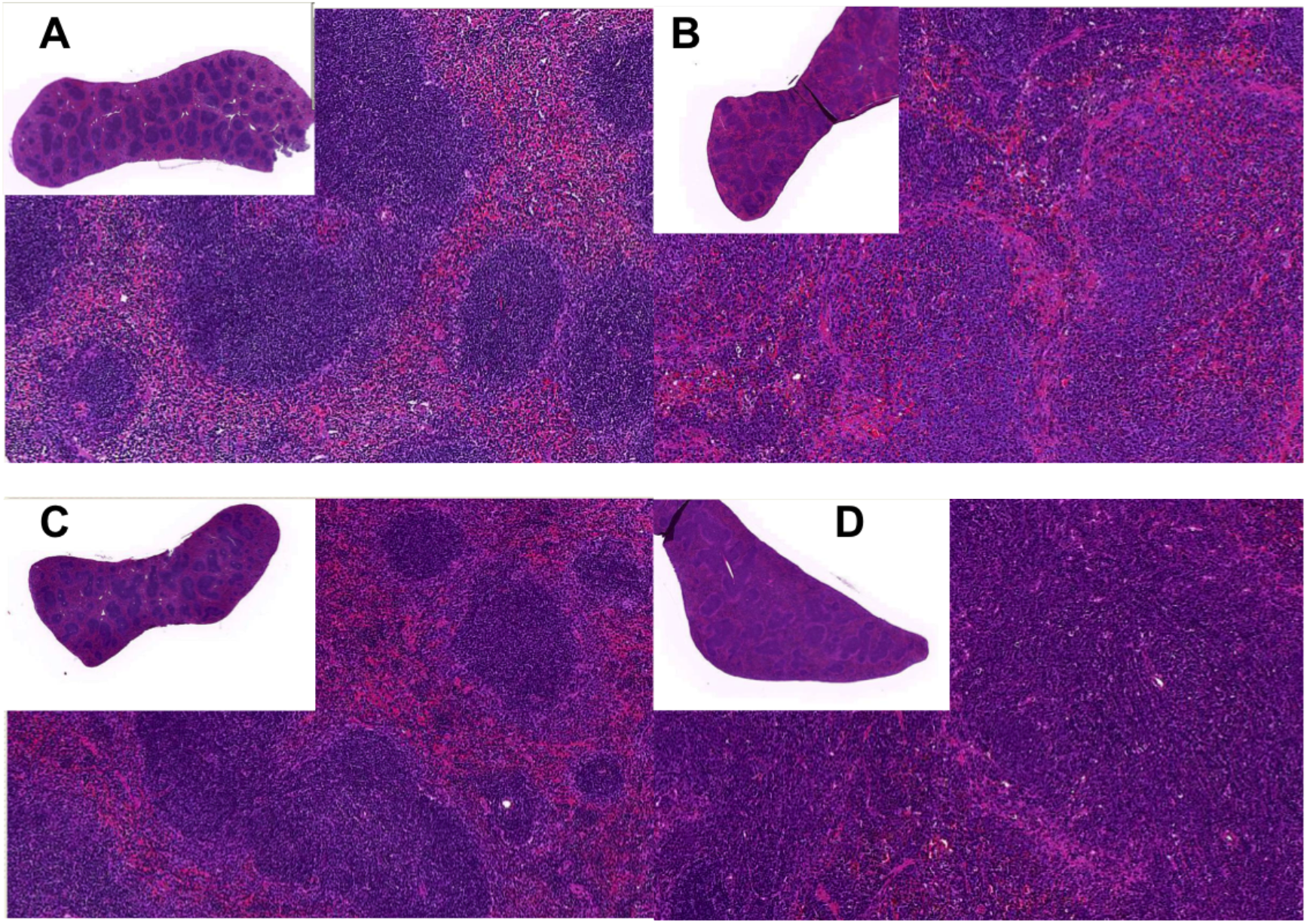
The M935K mutation causes an abnormal WBC response to infection. Representative H&E stained spleen sections from **(A)** uninfected WT, **(B)** infected WT, **(C)** uninfected Kcc1^M935K/M935K^, and **(D)** infected Kcc1^M935K/M935K^ mice. Sections are at 5× magnification; insets at 0.625× magnification.

**Figure S2:**
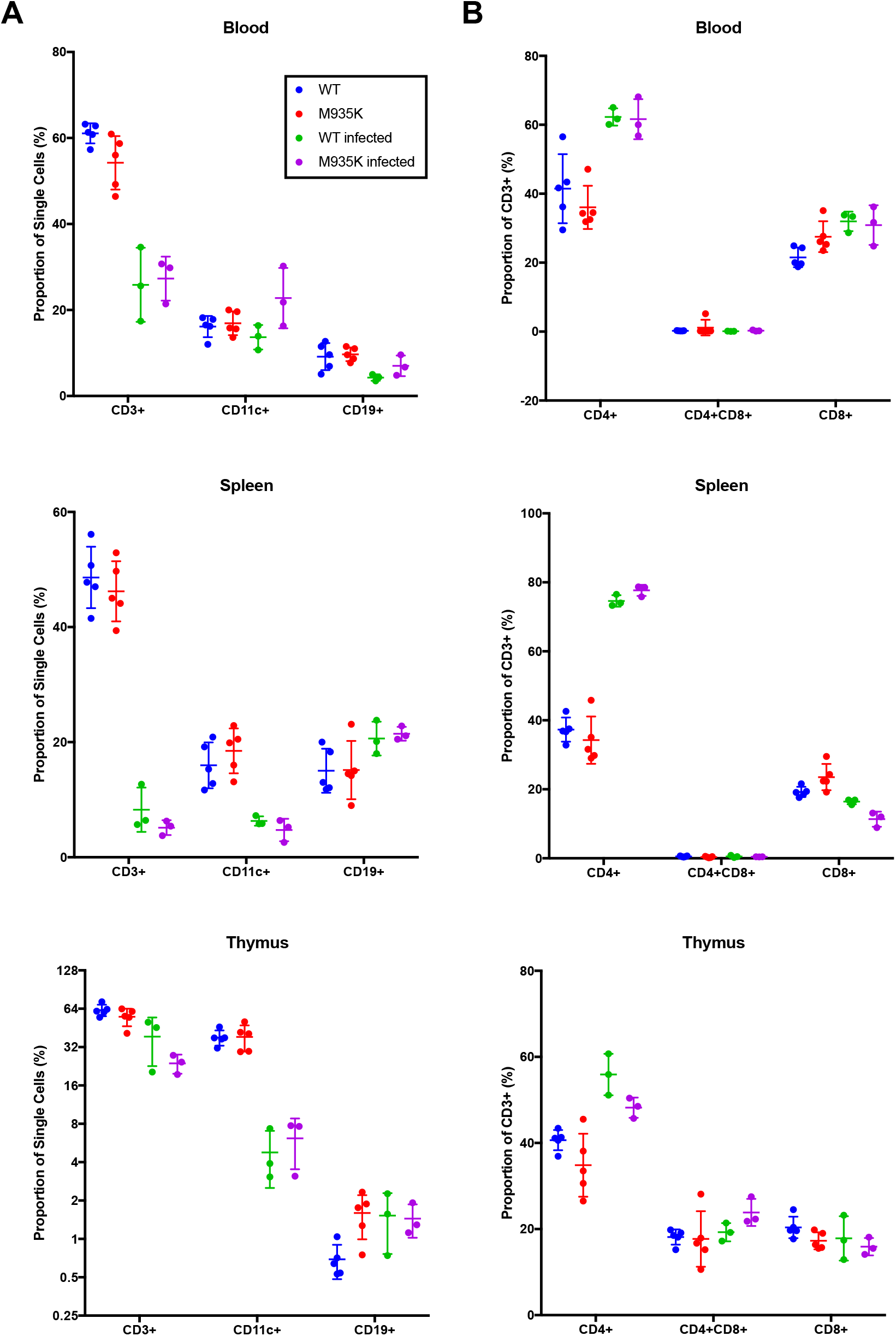
The Kcc1^M935K^ mutation does not alter immune cell populations in the blood, spleen, or thymus. **(A)** Average ± SEM proportion of lymphocytes that are CD3+, CD11c+, and CD19+ in the blood spleen and thymus. **(B)** Average ± SEM proportion of CD3+ cells that are CD4+, CD8+, and CD4+CD8+ in the blood spleen and thymus. Black circles = uninfected WT (n=5), grey circles = uninfected Kcc1^M935K/M935K^ (n=5), black open squared = infected WT (n=3), grey open squares = infected Kcc1^M935K/M935K^ (n=3).

**Figure S3:**
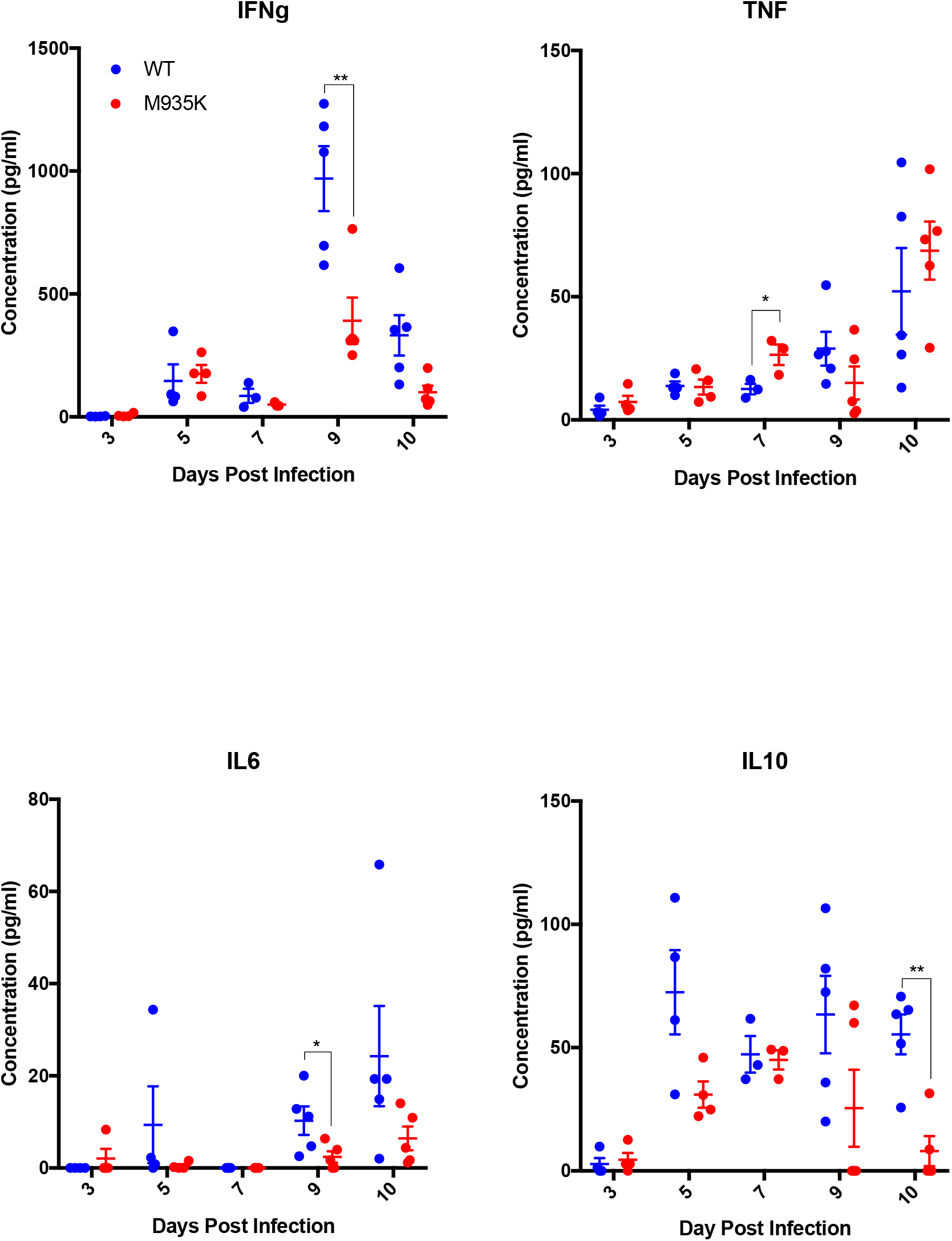
The Kcc1^M935K^ mutation alters the inflammatory response to *P. berghei* infection. Average ± SEM concentration of cytokines in the plasma of WT and Kcc1^M935K/M935K^ mice during infection with *P. berghei*. **P<0.01, Significance calculated using unpaired t test

